# *Psidium guajava* derived carbon nanoparticles: A promising red emissive cellular bioimaging agent

**DOI:** 10.1101/2023.03.20.533411

**Authors:** Shivani Mehta, Udisha Singh, Ketki Barve, Dhiraj Bhatia

## Abstract

We report a simple, cost-effective, microwave-assisted green synthesis route of red-emitting fluorescence carbon nanoparticles (CNPs) using *Psidium guajava* (Guava leaves). The synthesis of CNPs is a simple, affordable, and rapid method of producing carbon nanoparticles. The CNPs were characterized by various spectroscopic and microscopic techniques. Atomic force microscopy studies showed that the average size of CNPs is approximately 50 nm. The CNPs exhibited excellent photoluminescence properties with a maximum emission at 677 nm, making them suitable for bioimaging applications. The Ionic, photostability, and thermal stability of CNPs were also checked to understand their robustness. Retinal pigment epithelium (RPE) cells were exposed to these nanoparticles and showed very efficient uptake, some fraction of it also getting targeted to the nucleus, indicating that CNPs are non-toxic and biocompatible for future biological experiments. The results indicate that guava leaves can be a promising source for the synthesis of red emissive CNPs through the very simple method of synthesis and with bioimaging applications.

## 1. Introduction

Many precursor materials are being used for the synthesis of fluorescent carbon nanoparticles (CNPs), but among them, green synthesis of CNPs gain much attention as they are cost-effective and environmentally friendly. The development of such cost-effective, sustainable, and tuneable fluorescent CNPs will broaden the application of CNPs in bioimaging, sensing, drug delivery, catalysis, photovoltaics, and more. The most characteristic property of CNPs is fluorescence properties under ultraviolet (UV) irradiation. Most of the CNPs show fluorescence in the blue-green region. Very few CNPs green synthesized show fluorescence in the near-infrared region^1^. The CNPs we have synthesized show fluorescence in the red region of electromagnetic spectra. These red-emitting carbon nanoparticles (CNPs) were synthesized via microwave-assisted methods using Psidium guajava (Guava) leaves as a green synthesis route. This approach offers a facile and costeffective strategy for producing CNPs from biomass as a carbon source, as it is both environmentally friendly and naturally accessible when compared to alternative carbon sources^2^. The use of this natural biomass as a low-value resource can convert it into a valuable material, offering advantages such as high availability, low cost, eco-friendly, and renewability^3^. Numerous researchers have investigated the green synthesis of carbon nanoparticles (CNPs) utilizing plant resources as a precursor, instead of using chemical compounds, due to the sustainability of such an approach and the presence of heteroatoms, such as nitrogen, sulfur, boron, and phosphorus^3^. The presence of chlorophyll within green leaves, which possess porphyrin structures, results in substantial near-infrared (NIR) absorption and emission. However, due to its hydrophobicity and instability, porphyrin cannot be directly employed as a fluorescent probe in the field of bio-imaging^4^. The majority of the synthesized CNPs described in the literature exhibit a blue-to-green fluorescence that is toxic to living things including cells and biosystems^4^. Blue to green fluorescence carbon nanoparticles have been found to generate reactive oxygen species (ROS) which are molecules that can cause oxidative stress and damage to cells^5^. Consequently, the development of red or NIR-light-emitting CNPs is crucial^4^. Very limited studies have so far successfully developed red/NIR light-emitting CNPs. For example, Tseng et al. and his coworkers prepared CDs (630 nm) by using hydroxyl-radical-induced degradation of graphene oxide^6^. Rajita Ramanarayanan et al. used guava leaves for the synthesis of carbon dots through the hydrothermal method with water and ethanol as a solvent; they prepared carbon dot-TiO2 nanocomposite having high photocatalytic activity and the ability to degrade methylene blue dye^7^. Bharmore et al. synthesize fluorescent CNPs without any surface passivation from *Pyrus pyrifolia* (pear) fruit through a one-step hydrothermal method and applied in bioimaging in *bacillus subtilis* bacterial cell, and also used as a biosensor^8^. Most synthesis methods typically employ temperatures of approximately 150°C or higher, with the most commonly used techniques being hydrothermal or reflux methods. These methods generally require a timespan of more than one hour, whereas the microwave-assisted approach can be completed in less than half an hour.

Herein, we present a novel type of CNPs that emits red fluorescence through microwave-assisted method. Because it is a simple, cost-effective, and quick approach to synthesizing carbon nanoparticles. It contains microwave irradiation with a wide range of wavelengths from 1mm to 1m ^9^. Due to this, microwaves provide homogeneous heating to the precursor and break the chemical bond that exists in the precursor. The microwave approach is currently replacing the use of the traditional method since it is more eco-friendly, uses less energy and heats up quickly, and doesn’t require complex machinery. Overall, it accelerates the reactions and improves the quality of nanoparticles^10^. Overall, the CNPs we have synthesized show promising results and they can be further modified to be used as a good bioimaging tool in the future.

**Scheme 1.**
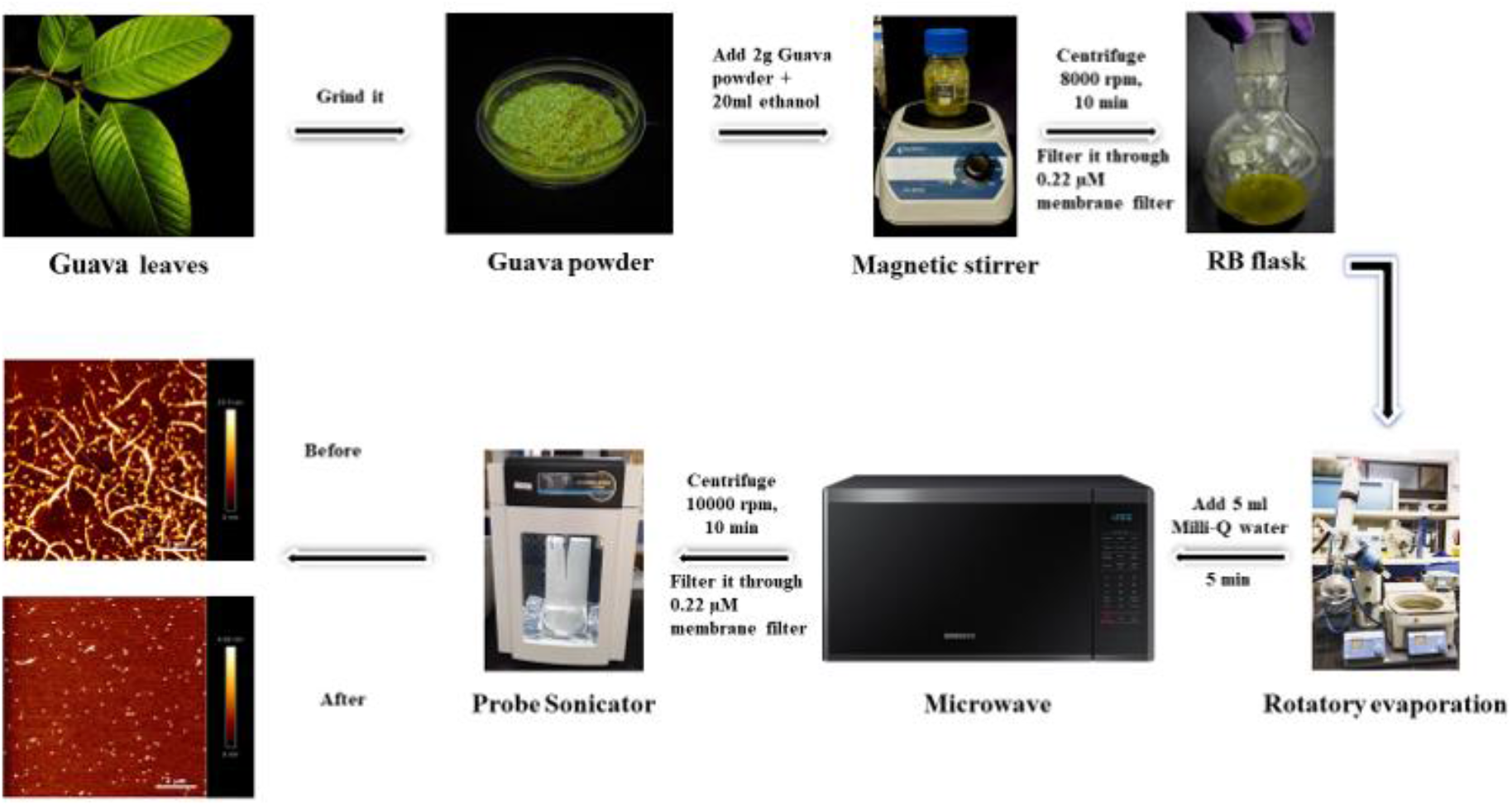
Schematic representation of synthesized CNPs. Red emitting carbon nanoparticles are synthesized through microwave-assisted methods by using Guava leaves. The Guava leaves were first dried under the sun and then ground to obtain powder form. The power obtained is dispersed in 20 ml ethanol and stirred on a magnetic stirrer. The solvent is then evaporated using rotary evaporator to get a slurry. The slurry obtained is dispersed in water and undergoes microwave treatment. The extract obtained after treatment is dispersed in ethanol and filtered through a syringe filter, then probe sonicated. The final CNP powder was obtained after the extract evaporated again in a rotary evaporator.

## 2. Results and Discussions

### 2.1 Synthesis and characterization of CNPs

Red emissive carbon nanoparticles are synthesized through the microwave-assisted method and facile bottom-up method which is a widely used green synthesize approach. In detail, 2 g psidium guava powder dispersed in 20 ml ethanol is kept in a magnetic stirrer for 4 hr and centrifuged for 10 min at 10000 rpm. Ethanol is evaporated in rotary evaporation until the slurry is obtained. Add 5 ml Milli Q water and microwave it for 5 min. The extract obtained was dispersed in ethanol and filtered out through a 0.22 µm membrane filter. The extract was then a probe sonicator for 30 min. Rotary evaporation evaporates the ethanol to obtain a powder. The powder was dispersed into ethanol and the same procedure was carried out but the powder was dispersed in water instead of ethanol. The obtained powder was then used for further characterization and experimental purposes. To determine the size and morphology, we utilized transmission electron microscopy (TEM). The TEM images indicated that the particles were evenly distributed and possessed a spherical disk-like structure. The average diameter of the fluorescent CNPs, as determined by counting 50 nanoparticles, was approximately 23.29 ± 2.7 nm, displayed in **figure 1a,b**. The measurement of particle size was conducted using ImageJ software with a scaled image, and the data were analyzed using Origin software. We obtained a characteristic halo ring SAED diffraction pattern of CNPs, which proves that they are amorphous. The shape and size were further validated by the atomic force microscopy (AFM) image (**figure 1c**).

**Figure 1.**
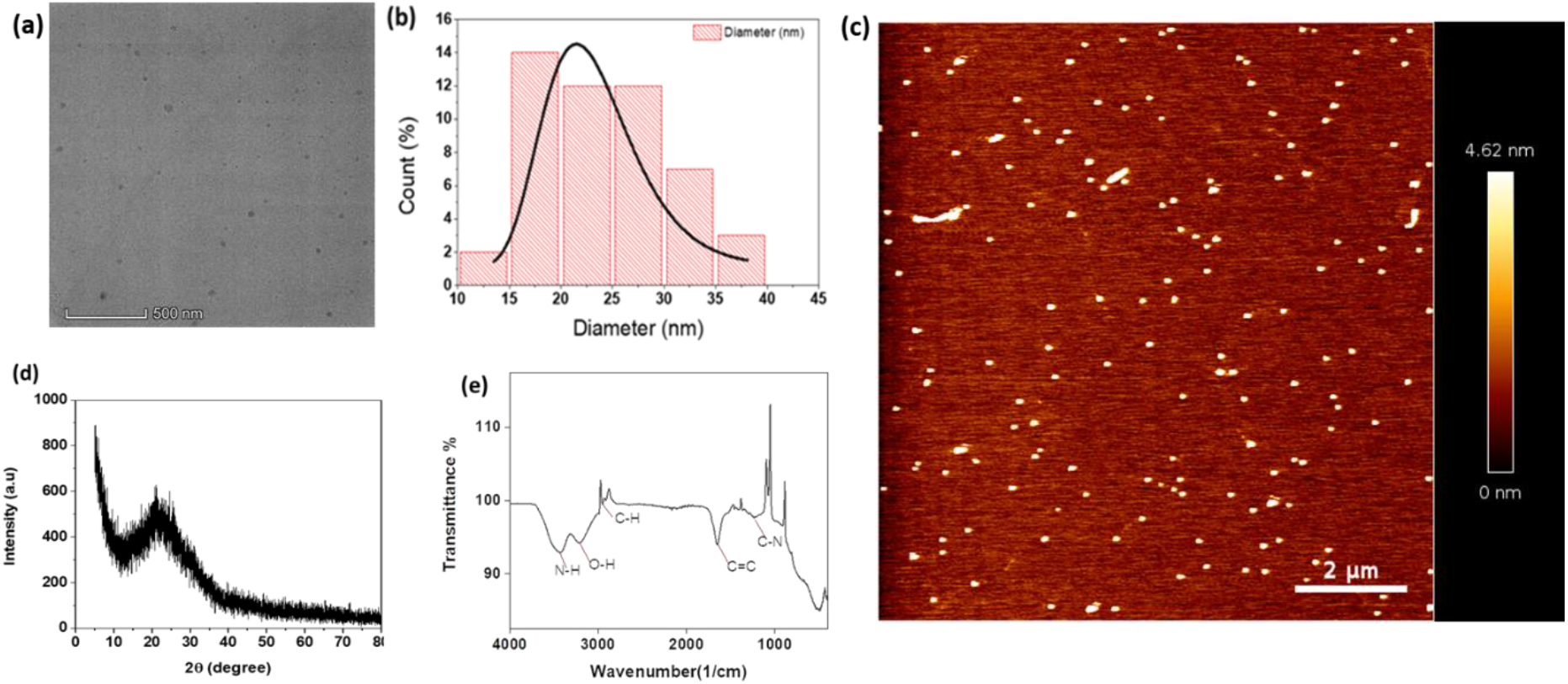
Characterization of red fluorescent CNPs. (**a,b**) Transmission electron microscopy (TEM) images of CNPs show spherical-shaped CNPs with an average particle size of 23.29 ± 2.7 nm. The scalebar is set at 500 nm (**c**) Atomic force microscopy (AFM) image of CNPs showing the similar sized CNPs with the scalebar set at 2µm (**d**) X-Ray diffraction (XRD) spectra of CNPs The broad peak at 23° shows that CNPs are amorphous in structure. The 2ϴ range for CNPs was kept between 10° to 80° (**e**). The functional groups attached to the surface of CNPs are confirmed by Fourier transform infrared(FTIR) spectroscopy. 0.1mg/ml of CNPs were drop cast on FTIR detection mode to get readings.

We conducted X-ray diffraction (XRD), and Fourier transform infrared (FT-IR) spectroscopy investigations to better understand the structure and composition of the fluorescent CNPs. Evidence for the generation of amorphous graphitic carbon having sp2 hybridization may be seen in the wide diffraction peak detected at about 23°C, which correlates to the (100) plane of carbon, as shown in **figure 1d**. The Fourier transform infrared spectrum is utilized to identify the functional groups that exist on the surface of nanoparticles. In the case of CNPs, chemical bonds are formed between N, H, C, and O elements. We obtained N-H stretching (3432 cm^-1^), O-H stretching (3206 cm^-1^), C-H stretching (2948 cm^-1^), C=C stretching (1646 cm^-1^), and C-N stretching (1227 cm^-1^). The absorption peak obtained at (3432 cm^-1^) confirms that the presence of primary amine on the surface of CNPs, (3206 cm^-1^) means (O-H stretch) ensures the presence of the carboxylic group, (2948 cm^-1^) and (1646 cm^-1^) confirms that presence of alkene group, (1227 cm^-1^) confirms that presence of amine group on the surface of CNPs as shown in **figure 1e**. The presence of the C-N bond on the surface of CNPs proves that amines are present as functional groups on the surface of CNPs, which proves heteroatom doping of CNPs. The presence of heteroatoms in functional groups can shift the emission spectra of CNPs from blue to red regions of the electromagnetic spectrum. According to reports, the existence of the -OH group in CNPs has demonstrated bioactive and chemically active traits that diminish the impact of cellular redox activity^11^.

### 2.2 Optical properties of CNPs

To comprehend the optical characteristics of CNPs, we analyzed their UV-VIS absorption spectra and fluorescence emission spectra. The stock solution prepared was 22mg/ml of CNPs and then further diluted to get the absorption spectra of CNPs. The maximum absorption peak showed in **figure 2a** at 277 nm and the tail expanded in the visible region due to the transition of electron n-π* of the C=O bond. The fluorescence (FL) spectra showed emission at different excitation wavelengths. The highest peak of emission of CNPs was obtained at 677 nm at an excitation wavelength of 280 nm. The fluorescence intensity is 4095 a.u at 677 nm, as shown in **figure 2b**. The emission wavelength is in the range of 650 nm to 750 nm. The CNPs showed green color under daylight and red color under UV light. The CNPs show the excitation wavelength-independent fluorescence spectra. With an increase in excitation wavelength, there was no shift in the emission spectra.

**Figure 2.**
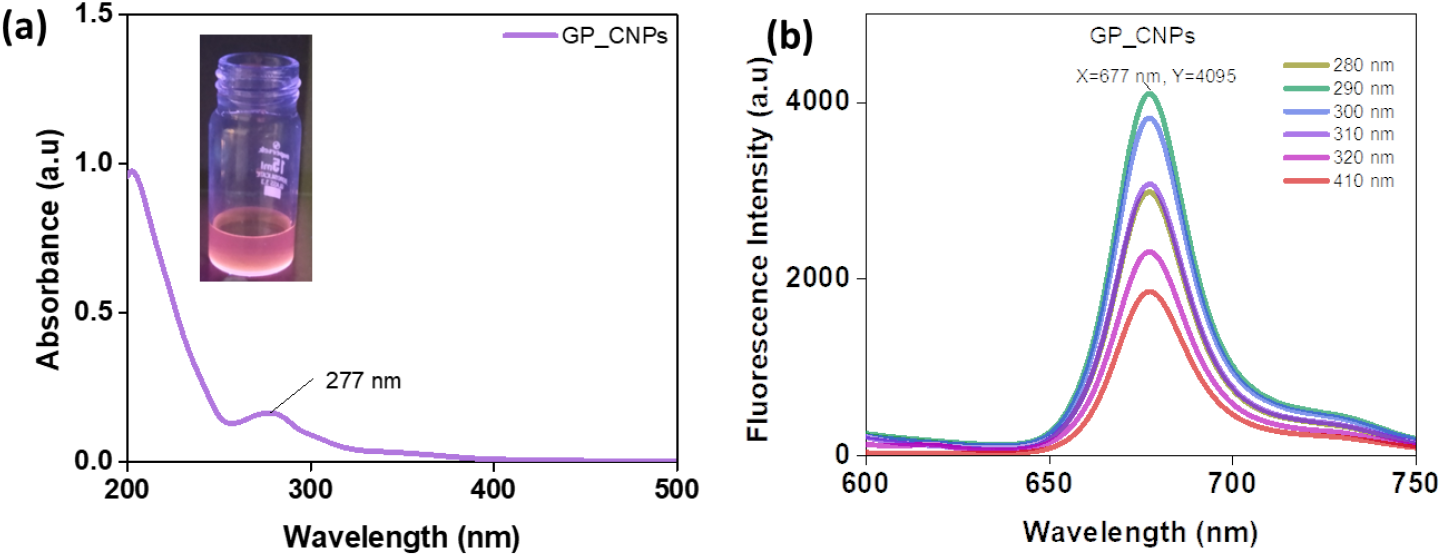
(**a**) The graph demonstrating CNPs’ ultraviolet-visible spectra with absorbance at 277 nm (**b**) Emission spectra of CNPs at 280 nm excitation revealed λ^max^ 677 nm emission wavelength.

We further investigated how different pH levels affect the fluorescence of CNPs. The CNPs (22 mg/mL) dispersed in MilliQ water where pH was adjusted to 2, 4, 5, 7, 9, and 12, and their fluorescence spectra were studied. As the pH increased, the fluorescent CNPs exhibited a decrease in fluorescence intensity. The highest fluorescence intensity peak was observed at pH 4, while the lowest fluorescence intensity peak was observed at pH 12, as shown in **figure 3a**. Between pH 4 and pH 7, fluorescence intensity decreased by around 24%, whereas between pH 7 and pH 12, it decreased by 6%. The majority of CNPs are pH sensitive due to the functional groups that are present on their surface being protonated and deprotonated, which changes the optical properties of CNPs.

**Figure 3.**
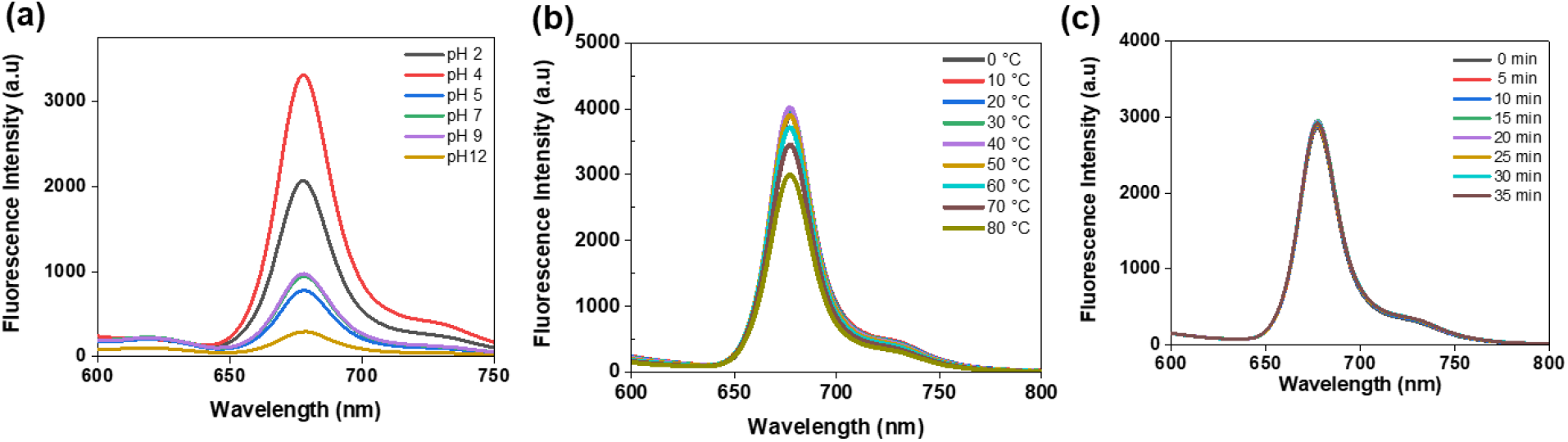
(a) Fluorescence spectra of CNPs at a range of 2 - 12 pH. (b) Fluorescence spectra at different temperatures (0°C – 80°C). (c) Fluorescence spectra were captured from 0 to 35 min under constant irradiation of UV light. 30 µg/ml of CNP dispersed in water and used for the fluorescence spectroscopic analysis to check the ionic, thermal and photostability of CNPs.

**Figure 4.**
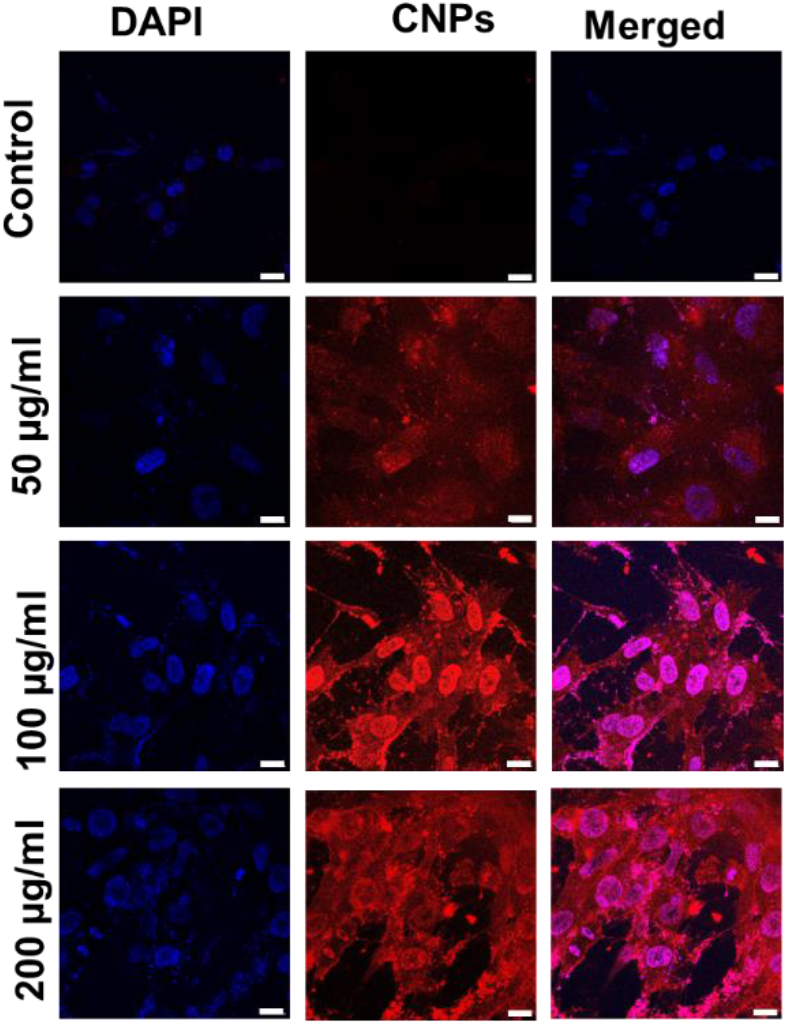
Showing the confocal images of CNPs treated Retinal Pigment epithelial (RPE**-**1) cells. Concentration-dependent studies of CNPs were done. Control is not treated with CNP and hence shows no fluorescence when excited at 633 nm excitation laser. The fluorescence intensity keeps on the increase with the increase in the concentration of CNPs. The scale bar for confocal images is maintained at 10 µm.

The thermal stability of CNPs was further investigated by subjecting them to a range of temperatures from 10°C to 80°C while monitoring changes in their fluorescence spectra using a spectrophotometer. Initially, the fluorescence intensity was obtained at the same peak shown in **figure 3b** from 10°C to 50°C. The decrease in fluorescence intensity from 60°C to 80°C is approximately 7%. So, here we conclude that fluorescence intensity is almost constant but is slightly decreased after 60°C because of electronic vibrations in the presence of thermal energy leading to non-radiative energy release^12^. Further, we studied the time-dependent fluorescence intensity in the 5 min time interval for 30 min, but the intensity was stable by the time. The peak was observed at the same intensity shown in figure **3c**. Both temperature and continuous light-dependent studies prove that the red-emitting CNPs we have synthesized are quite thermo and photostable.

### 2.3 Cellular uptake of carbon nanoparticles

To study the bioimaging applications of the CNPs, confocal microscopy studies were done on the mammalian retinal pigmental epithelial cells (RPE1) incubated with QDs at different concentrations. Concentration-dependent studies revealed that when cells were treated with varying concentrations of CNPs from 50, 100, and 200 g/ml for 30 min at 37ºC, the CNPs were efficiently uptaken within cells and showed intracellular fluorescence, sometimes even from the nucleus; and the cellular fluorescence intensity increased with the concentration of CNPs. The CNPs emission from within the cells was obtained in red channel and at 633 nm laser excitation and the fluorescence signal increased with increase in concentration of CNPs.

## 3. Conclusions

We demonstrate the synthesis of red fluorescent carbon nanoparticles of size ranging between 10 to 40 nm through the microwave-assisted method by using *Psidium guajava* (Guava leaves) as precursor material and ethanol as a solvent. Red emissive CNPs have great potential for bioimaging. To understand the physical and chemical properties of CNPs various characterization techniques were employed. The fluorescence intensity is dependent on the different solvents used and the CNPs are more soluble in ethanol as compared to water. pH also affects the fluorescence The fluorescence intensity is also dependent on pH and in inverse correlation, that is, when pH is increased, the intensity decreases. Furthermore, CNPs also show stability at high temperatures and continuous light exposure, which makes the CNPs quite thermostable and photostable. These CNPs are biocompatible and have been used for bioimaging in mammalian cells. The cellular uptake and intensity of fluorescence increases with an increase in the concentration of CNPs. Although very preliminary at this stage, the present work provides us with a prominent fluorescent material that can be used and can be further modified to enhance the optical properties of CNPs and make them more target oriented and functionalized for specific biological applications like bioimaging and delivery in future.

## 4. Material and Method

### 4.1 Material

The fresh guava leaves (*Psidium guajava)* were collected from the garden of IIT Gandhinagar, Gujarat, India. The experiment used deionized water with a conductivity of 18.2 M.CM@25°C. Ethanol absolute (Analytical CSS reagent). Syringe filter procured from Merck. Dulbecco’s modified Eagle’s medium (DMEM), penicillin−streptomycin, trypsin−EDTA (0.25%), and fetal bovine serum (FBS) from Gibco. No additional sterilization or treatment is required because all the substances were of excellent scientific grade.

### 4.2 Method to Synthesize red emissive CNPs

The microwave-assisted method was used to produce CNPs that emit bright red fluorescence. The fresh leaves of *Psidium guajava* (Guava leaves) were taken and washed with Milli-Q water to remove dust, then left to dry for two days at room temperature. The dried leaves are ground into powder in a mixer, and 2 grams of leaf powder added to 20 ml of ethanol are stirred using a magnetic stirrer. The extract was then centrifuged for 10 minutes at 8000 RPM. Filter the supernatant with a syringe filter (0.22m); then, rotary evaporation is used to evaporate the ethanol until a slurry is obtained. Add 5 ml milli-Q water, and microwave for 5 minutes. The extract is dispersed into 5 ml ethanol and centrifuged at 10,000 rpm for 10 min. Collect the supernatant and further filter it out through a syringe filter (0.22 µm). The extract was then a probe sonicator for 30 min. Again, rotational evaporation evaporates the ethanol to obtain a powder. Initially, the powder was dispersed in ethanol, and then the same procedure was followed, but the powder was dispersed in water instead of ethanol.

### 4.3 Various analytical methods used for the characterization of CNPs

FEI Titan Themis transmission electron microscope (TEM) (**300 kV**) was used to analyze the morphology of fluorescent CNPs. Brucker Nano wizard Sense atomic force microscope (AFM) was used to capture both two-dimensional (2D) and three-dimensional (3D) imaging of CNPs.The X-ray diffraction (Rikagu smart lab 9KW (XRD)) method was employed for phase identification and to analyze the structural characteristics. FP8300 Jasco spectrofluorometer (Japan) was used to obtain the fluorescence spectra within the range of 280 to 410 nm. Spectro Cord-210 Plus Analytik Jena (Germany) was employed to measure the UV-vis absorbance spectra. An FTIR spectrometer from Spectrum 2 by PerkinElmer was used to capture the Fourier transform infrared (FT-IR) spectra of CNPs in the attenuated total reflectance (ATR) mode. The scanning was conducted over the range of 400 cm^-1 to 4000 cm^-1. Using a Leica TCS SP8 confocal laser scanning microscope with a resolution of 63x, fixed sample cell imaging was carried out.

### 4.4 Cell culture studies

The retinal pigment epithelial (RPE) cell line used in this experiment was gifted from Prof. Ludger Johannes, Curie Institute, Paris. DMEM complete media was used for growing cells, and all the experiments were carried out using 1X PBS at a pH of 7.4.

#### 4.4.1 Bioimaging studies using confocal microscopy

Cellular studies were performed to study the bioimaging properties of CNPs using mammalian cell lines. RPE-1 cells were seeded on coverslips in a 4-well plate with a density of approximately 20,000 cells/well, treated with CNPs, and incubated at 37°C for a specific duration. After treatment, the cells were fixed using 4% PFA at 37°C for 15 minutes, followed by washing with PBS thrice to eliminate any residual PFA or impurities for accurate imaging under a microscope. Hoechst dye was used to stain the nuclei, and the fixed cells were imaged using a confocal laser scanning microscope at a resolution of 63X. The obtained images were quantified using Fiji ImageJ software.

## Acknowledgements

We sincerely thank all the members of the DB group for critically reading the manuscript and for their valuable feedback. US thanks IITGN-MHRD, GoI for the PhD fellowship. KB thanks Dr D Y Patil Vidyapeeth, Pune, for permission for her MTech internship at IITGN. SM thanks Sardar Patel University for permission to carry out research at IITGN. DB thanks SERB, GoI for Ramanujan Fellowship, IITGN for the startup grant, and DBT-EMR, Gujcost-DST & GSBTM for research grants. Imaging facilities of CIF at IIT Gandhinagar are acknowledged.

## References

(1) Chahal, S.; Macairan, J.-R.; Yousefi, N.; Tufenkji, N.; Naccache, R. Green Synthesis of Carbon Dots and Their Applications. RSC Adv. 2021, 11 (41), 25354–25363. https://doi.org/10.1039/D1RA04718C.

(2) Meng, W.; Bai, X.; Wang, B.; Liu, Z.; Lu, S.; Yang, B. Biomass-Derived Carbon Dots and Their Applications. ENERGY Environ. Mater. 2019, 2 (3), 172–192. https://doi.org/10.1002/eem2.12038.

(3) Kanwal, A.; Bibi, N.; Hyder, S.; Muhammad, A.; Ren, H.; Liu, J.; Lei, Z. Recent Advances in Green Carbon Dots (2015–2022): Synthesis, Metal Ion Sensing, and Biological Applications. Beilstein J. Nanotechnol. 2022, 13 (1), 1068–1107. https://doi.org/10.3762/bjnano.13.93.

(4) Li, L.; Zhang, R.; Lu, C.; Sun, J.; Wang, L.; Qu, B.; Li, T.; Liu, Y.; Li, S. In Situ Synthesis of NIR-Light Emitting Carbon Dots Derived from Spinach for Bio-Imaging Applications. J. Mater. Chem. B 2017, 5 (35), 7328–7334. https://doi.org/10.1039/C7TB00634A.

(5) Singh, U.; Shah, K.; Kansara, K.; Kumar, A.; Bhatia, D. Novel Class of Yellow Emitting Carbon Dots Stimulate Collective Cell Migration and 3D Uptake in Vivo. bioRxiv July 4, 2022, p 2022.07.04.498723. https://doi.org/10.1101/2022.07.04.498723.

(6) Ke, C.-C.; Yang, Y.-C.; Tseng, W.-L. Graphene Quantum Dots: Synthesis of Blue-, Green-, Yellow-, and Red-Emitting Graphene-Quantum-Dot-Based Nanomaterials with Excitation-Independent Emission (Part. Part. Syst. Charact. 3/2016). Part. Part. Syst. Charact. 2016, 33 (3), 121–121. https://doi.org/10.1002/ppsc.201670008.

(7) Ramanarayanan, R.; Swaminathan, S. Synthesis and Characterisation of Green Luminescent Carbon Dots from Guava Leaf Extract. Mater. Today Proc. 2020, 33, 2223–2227. https://doi.org/10.1016/j.matpr.2020.03.805.

(8) Bhamore, J. R.,,,, Jha, S.,,,, Singhal, R. K.,,,, Park, T. J.,,,, Kailasa, S. K. Facile Green Synthesis of Carbon Dots from Pyrus Pyrifolia Fruit for Assaying of Al3+ Ion via Chelation Enhanced Fluorescence Mechanism. J. Mol. Liq. 2018, 264, 9–16. https://doi.org/10.1016/j.molliq.2018.05.041.

(9) Khayal, A.; Dawane, V.; Amin, M. A.; Tirth, V.; Yadav, V. K.; Algahtani, A.; Khan, S. H.; Islam, S.; Yadav, K. K.; Jeon, B.-H. Advances in the Methods for the Synthesis of Carbon Dots and Their Emerging Applications. Polymers 2021, 13 (18), 3190. https://doi.org/10.3390/polym13183190.

(10) Schwenke, A. M.; Hoeppener, S.; Schubert, U. S. Synthesis and Modification of Carbon Nanomaterials Utilizing Microwave Heating. Adv. Mater. 2015, 27 (28), 4113–4141. https://doi.org/10.1002/adma.201500472.

(11) Lu, F.; Yang, S.; Song, Y.; Zhai, C.; Wang, Q.; Ding, G.; Kang, Z. Hydroxyl Functionalized Carbon Dots with Strong Radical Scavenging Ability Promote Cell Proliferation. Mater. Res. Express 2019, 6 (6), 065030. https://doi.org/10.1088/2053-1591/ab0c55.

(12) Lakowicz, J. R. Introduction to Fluorescence. In Principles of Fluorescence Spectroscopy; Lakowicz, J. R., Ed.; Springer US: Boston, MA, 1999; pp 1–23. https://doi.org/10.1007/978-1-4757-3061-6_1.

